# Understanding Vascular Endothelial Cell Behavior Using a Mechanical Strain Gradient Generated by an Electromagnetic Stretching Device

**DOI:** 10.1101/2022.10.20.513030

**Authors:** Michelle L. Yang, Catalina Zuo, Jiafeng Liu, Kun Man, Zhenyu Zuo, Yong Yang

## Abstract

Cardiovascular diseases cause an estimated 17.9 million deaths globally each year (World Health Organization). Endothelial cells that line the vasculature and the endocardium are subjected to cyclic mechanical stretch. Deviation from physiological stretch can alter the endothelial function, having the risk of atherosclerosis and myocardial infarction. To understand the mechanical stretch effects, cell culture platforms that provide mechanical stretch have been developed. However, most of them have fixed strain and frequency, sometime not in the pathophysiological range. We thus developed a novel, electromagnetically driven, uniaxial stretching device, where cells were grown on a flexible polydimethylsiloxane (PDMS) membrane mounted onto a 3-D printed track. The strain of the membrane was readily controlled by tailoring the track design and the frequency was determined by electromagnetic actuation. Furthermore, the mechanical strain gradient was generated on a PDMS membrane with a tapered thickness. This strain gradient, ranging from 1.5% to 40%, covered both physiological and pathological vascular stretch ranges. When human vascular endothelial cells were subjected to the cyclic stretch, the cells exhibited strain-dependent cell and nuclear orientation and elongation perpendicular to the stretching direction, compared to the random cell and nuclear orientation under the static condition. However, the overstretching led to deviation from the aforementioned orientation and elongation, and impaired the tight junctions, leading to a leaky endothelium. This novel, versatile, cost-effective, pathophysiologically relevant stretching device provides a useful platform for advancement of vascular disease research and treatment.

## 1. Introduction

Mechanical stretch plays a critical role in the human body with its importance to the function and homeostasis of cells, tissues, and organs. For example, endothelial cells lining the vascular system undergo physiological cyclic mechanical deformation due to pulsatile blood pressure and circulating blood flow through arteries, veins, and capillaries. Deviation from physiological mechanical stretch can alter the endothelial cell phenotype and function, which can lead to pathological changes and result in cardiovascular diseases such as atherosclerosis (buildup of plaque in arteries) and myocardial infarction (heart attack)^1^. Of note, the dysfunction of human cerebral microvascular endothelial cells (hCMECs) contributes to the onset and progression of neurodegenerative diseases such as Alzheimer’s and Parkinson’s diseases^2^. The physiological range of vascular mechanical strain *in vivo* is 5-10% and a strain above 20% is often seen in hypertension, leading to serious, often fatal health conditions^3^. In the cardiovascular system, hypertension forces the heart to work harder to pump blood throughout the body due to the thickening and hardening of arteries. It is thus essential to understand the role of mechanical stretch in the normal function of cells, tissue, and organs, and human disease progression.

Cell culture platforms that provide mechanical stretch, from one-dimensional (1-D) uniaxial stretch, 2-D biaxial or circumferential stretch, to 3-D radial stretch, have been developed. These devices have utilized different mechanisms of stretching a flexible membrane^4^, including mechanical actuation (via a motor)^5–6^, pneumatic actuation (via pressure with fluids)^7–8^, piezoelectric actuation (via electrical energy to generate mechanical displacement)^9–10^, electromagnetic actuation (via magnetic force)^11^, and optical actuation (via energy and light)^12^. However, most of these existing platforms are hindered by a fixed strain and frequency, non-uniform strain, or require direct physical contact with the cells, all unfit for the observation of cell behavior in the pathophysiological range. For example, the reported electromagnetically actuated device has multiple downfalls including a non-physiologically relevant magnitude and frequency of strain as well as non-uniform strain^11^. Therefore, there is a need for pathophysiologically relevant, effective *in vitro* models to study the role of mechanical stretch in cell regulation.

In the present work, we have developed a simple, cost-effective, electromagnetically driven 1-D stretching device with versatile strain, frequency, and potentially material properties. Notably, we were able to generate a strain gradient using a tapered polydimethylsiloxane (PDMS) membrane. Our computational and experimental characterization of the PDMS membrane confirmed the formation of strain gradients of pathophysiological relevance. Furthermore, we demonstrated that vascular endothelial cells, using hCMECs as model cells, grown on PDMS membrane in the device responded differently to the various strains along the gradient.

## 2. Materials and Methods

### 2.1. Fabrication of the electromagnetic stretching device

The stretching device consisted of a stretching part and an electromagnetic actuator. The stretching part was comprised of several components, including a track, clamps, a container, and multiple dowels (Figure 1). All these components were designed on SolidWorks (Dassault Systems SOLIDWORKS Co., Waltham, MA, USA) and printed with Elegoo Standard Photopolymer resin (Elegoo, Shenzhen, China) using a 3-D printer (Form 3, Formlabs, Somerville, MA, USA).

**Figure 1.**
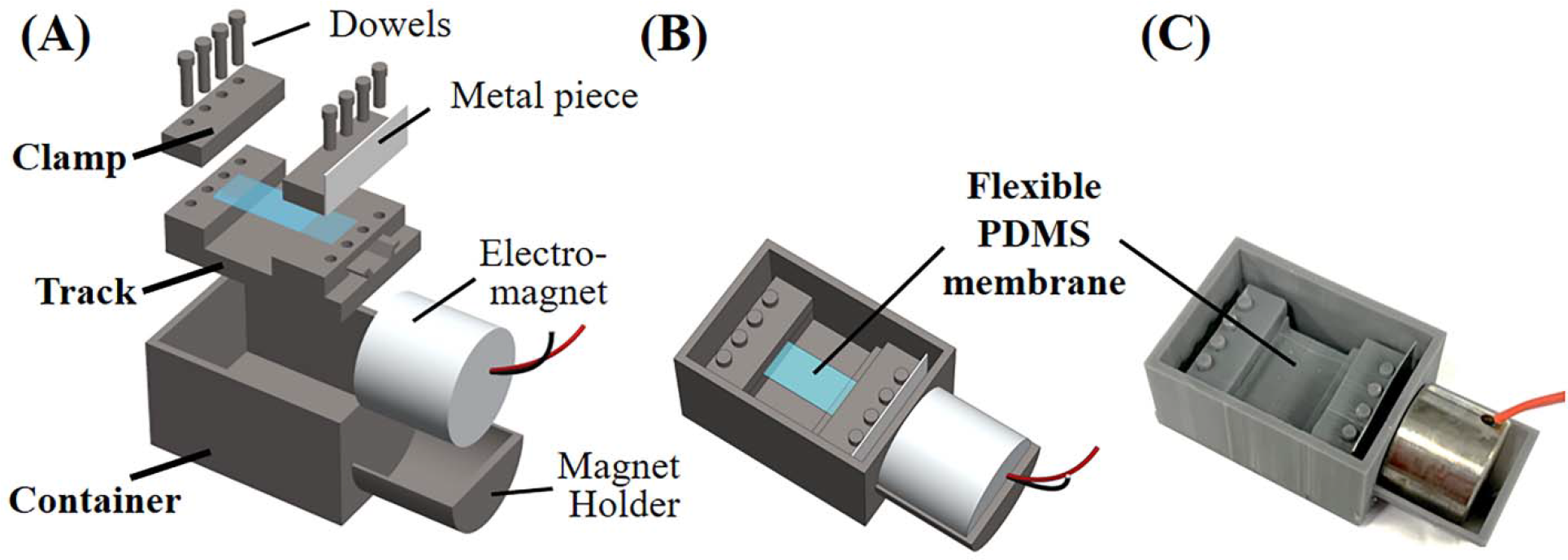
Design and photo of the electromagnetic stretching device. (A) The exploded view, (B) assembled view, and (C) a photo of the device.

### 2.2. Preparation of the stretchable membrane

The stretchable membrane was made of PDMS (Sylgard 184, Dow Corning, Midland, MI, USA). The PDMS membrane was prepared by spin-coating a mixture of PDMS resin and curing agent at a weight ratio of 10:1.05 on an overhead transparency plastic film, which was pre-trimmed into a rectangle, at 300 rpm for 60 seconds on a spin-coater (Laurell Technologies Co., North Wales, PA, USA). Next, a glass coverslip was placed on the PDMS mixture coated on the rectangular plastic film with one end touching the plastic film and another end elevated by two identical glass coverslips. In such a way, a PDMS membrane with a thickness gradient formed. After curing at 75 ◻ for 1 hour on a hotplate, the glass coverslip was carefully removed from the PDMS membrane and the PDMS membrane was then gently peeled off from the plastic film with a razor blade. The resulting membrane was trimmed using a razor blade into the dimensions of 1 cm × 3 cm (width × length) which had a thickness gradient of 300 μm on the thick end. When the PDMS membrane was mounted on to the stretching device between the fixed and moving components, 0.5 cm on each end was clamped between the device, and hence the actual PDMS membrane being stretched was 1 × 2 cm with a thickness gradient of approximately 250 μm to 50 μm.

### 2.3. SolidWorks simulation of the mechanical strain

3-D model for the gradient membrane was first built using SolidWorks to match the test membrane geometry. A custom material model for PDMS was then created in the SolidWorks simulation module. The material model was built with the mechanical and physical properties obtained from the literature^13^ and provided in Table 1. The linear elastic isotropic material model was used to define the linear strain behavior. The thinner side of the membrane was fixed and a 3 mm overall displacement was set as external load on the thicker side, corresponding to an overall strain of 15% on the membrane.

**Table 1.** Mechanical properties for PDMS

| Property | Value | Units |
| --- | --- | --- |
| Elastic Modulus | 1.685 | MPa |
| Poisson's Ratio | 0.499 | N/A |
| Shear Modulus | 0.56 | MPa |
| Mass Density | 965 | kg/m<sup>3</sup> |
| Tensile Strength | 5.69 | MPa |

### 2.4. Characterization of the mechanical strain

The strains experienced by the PDMS membrane were measured using a microscope (Evos XL Core, Thermo Fisher Scientific, Waltham, MA, USA) by examining the displacement of ink dots marked on the PDMS membrane. For the gradient membranes, three dots were drawn on the 1 cm × 2 cm PDMS membrane along the stretch gradient at 0.5 cm intervals starting from the thicker side of the membrane gradient (0.5 cm, 1.0 cm, 1.5 cm). The deformation of the dots during mechanical stretching were recorded using the microscope, and the strain at each of the points was quantified by analyzing the displacement of the dots using ImageJ (http://rsb.info.nih.gov/ij/index.html).

### 2.5. Cell Culture

Human cerebral microvascular endothelial cell line (hCMEC/D3; Cat#: CLU-512, Cedarlane, Burlington, NC, USA) were cultured in EBM-2 endothelial cell growth basal medium (Lonza, Bend, OR, USA) supplemented with EGM-2MV microvascular endothelial cell growth medium SingleQuots supplements (Lonza), 100 U/ml penicillin, and 100 μg/ml streptomycin (Life Technologies, Carlsbad, CA, USA).

The stretch device was oxygen plasma treated at the low power setting for 30 seconds in a plasma cleaner (Model PDC-001, Harrick Plasma, Ithaca, NY, USA) to change the chemical property of the PDMS membrane from hydrophobic to hydrophilic to facilitate cell adhesion. The PDMS membrane was then coated with 1 ml of 50 μg/ml rat collagen I (Corning, Corning, NY, USA) overnight in the incubator of 37 °C and 5% CO_2_. Cells were seeded at a density of 5 × 10^4^ cells/cm^2^ and cultured under the static condition for 4 hours, and then subjected to the mechanical stretch at an overall strain of 15% and a frequency of 1 Hz continuously for 16 hours. In comparison, the cells at 5 × 10^4^ cells/cm^2^ were cultured under the static condition for 20 hours as the control. The cells were then fixed for immunofluorescence staining.

### 2.6. Immunofluorescence staining

The cells on the membrane were fixed with 4% paraformaldehyde (PFA; Sigma-Aldrich, St Louis, MO, USA) for 30 minutes at room temperature, and blocked in a phosphate buffered saline (PBS; Thermo Fisher Scientific) solution containing 0.03 g/ml BSA, 1% goat serum (Gibco, Grand Island, NY, USA), and 0.2% Triton-X 100 (Sigma-Aldrich) for 1 hour. The samples were then incubated with Alexa Fluor 488 phalloidin (Life Technologies, 1:200 in PBS solution with 0.2% Triton-X 100) at 4 □ overnight. The nuclei were stained and mounted using ProLong Gold Antifade Reagent with DAPI (Life Technologies). The fluorescent images were taken by using a Nikon Ti Eclipse Fluorescence Microscope (Nikon, Melville, NY, USA).

### 2.7. Image analyses

The orientation and elongation of the nuclei were analyzed using ImageJ. The nucleus images were extracted by adjusting threshold of color to achieve best-fitted ellipses, from which the major and minor axis were determined. The orientation angle was defined as the angle between the nuclear major axis and the stretch direction. The elongation was defined by the ratio of nuclear major axis to minor axis and minus 1.

### 2.8. Statistical analysis

Statistical analysis was performed using GraphPad Prism software (GraphPad software, San Diego, CA, USA). P values between groups were calculated using unpaired two-tailed t-test. *p* < 0.05 was considered statistically significant.

## 3. Results

### 3.1. Design and fabrication of the electromagnetic stretching device

The electromagnetically driven 1-D stretching device encompassed a stretching part (moving component, fixed component, membrane, and track), a container, and an electromagnetic actuator as shown in Figure 1. The moving and fixed components were both connected to the track. The fixed component was connected to the base while the moving component was slid onto the track, with the ability to freely move with little friction acting upon it. The PDMS membrane was then secured on the device between the upper and lower pieces of the fixed and moving components as 0.5 cm of each end (both the thick and thin ends) of the flexible membrane was clamped down by conical dowels (Figure 2A and 2C). One end of the membrane was secured between the fixed component while the other end was secured between the moving component, allowing for uniform stretch as the moving component oscillated back and forth. As shown in Figure 2B, the moving component had a trapezoidal cutout slightly larger than the track, which had rounded edges and were shaped to reduce the amount of their contact area to reduce friction. As such, the moving component could move smoothly with a minimal amount of electromagnetic force to pull the PDMS membrane.

**Figure 2.**
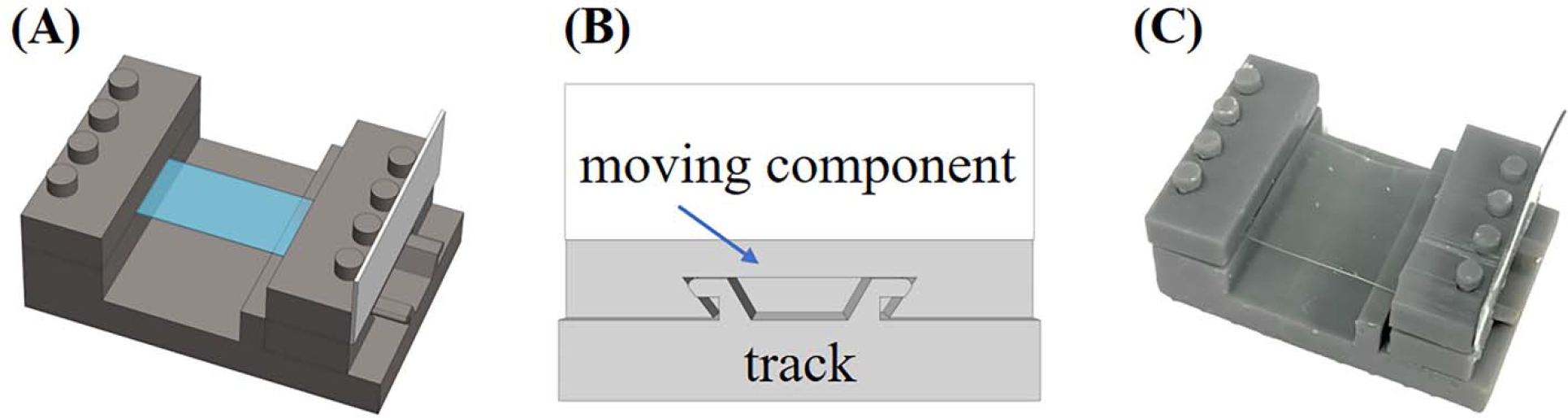
The uniaxial cell stretching part. (A) Illustration of the stretching part, (B) cross sectional view of the track with the moving component, and (C) the photo of the stretching part.

A rectangular piece of metal was attached onto the moving component as it was attracted by the electromagnet and therefore pulling the moving component towards the magnet with the disturbing force and being pulled back to the original position by the membrane’s restoring force, creating uniform oscillations. The membrane stretch was generated by an electromagnetic actuator that was built with an electromagnet (uxcell 5V 50N Electric Lifting Magnet Electromagnet; Amazon) controlled by an electric circuit containing a pulse width modulation (PWM) signal generator (ZK-PP2K pulse frequency generator; Amazon), a battery pack, and a switch (TWTADE Rocker switch; Amazon) (Figure 3). The PWM signal generator allowed the current to be turned on and off at a certain rate, and thus the frequency at which the magnet pulled could be controlled. In addition, the stretching part and cell culture medium were maintained within the container, with the magnet holder built to the outside of the container (Figure 1A). The separation of the electromagnetic actuator and the cell culture medium allowed for greater biocompatibility as viable cells were able to be maintained without exposure to contaminants. Note that magnet field strength is inversely proportional to the distance squared from the magnet. As a result, the magnetic field strength at the location of the metal decreases dramatically with distance. To maximize the amount of the magnetic force acting on the system, the wall between the electromagnet and the metal piece as thin as possible.

**Figure 3.**
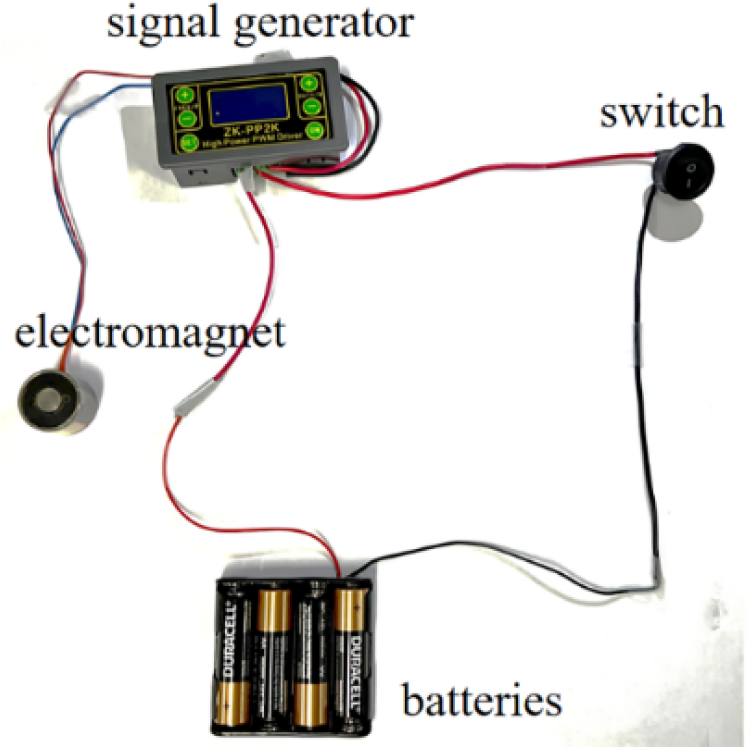
Electromagnetic actuator.

### 3.2. Generation of mechanical strain gradient

It was hypothesized that when the PDMS membrane with a gradient in thickness would result in varying strains at different locations. Along the stretch direction, the thicker areas of the membrane experienced a smaller strain than the thinner side of the membrane, which experienced greater strain.

To profile the strain gradient, the SolidWorks simulation was conducted. The dimensions of the membrane area stretched were 1 cm × 2 cm with a thickness gradient varying from 250 μm to 50 μm (Figure 4A). The material properties of PDMS were shown in Table 1. The simulation was based on an overall strain of 15%, which stretched the length from its original 2 cm to 2.3 cm. The SolidWorks simulation predicted a large range of strains up to 40% (Figure 4B). For instance, the theoretical strains at 0.5 cm, 1.0 cm, and 1.5 cm from the thicker end of the PDMS membrane were 7.4%, 10.0%, and 15.8%, respectively (Table 2). Note that the strain gradient was non-linear.

**Table 2.** Comparison of the SolidWorks simulation and the measured strains of the gradient

| Location from the thicker end | 0.5 cm | 1.0 cm | 1.5 cm |
| --- | --- | --- | --- |
| Observation | 9.8% | 16.0% | 21.0% |
| Simulation | 7.4% | 10.0% | 15.8% |

**Figure 4.**
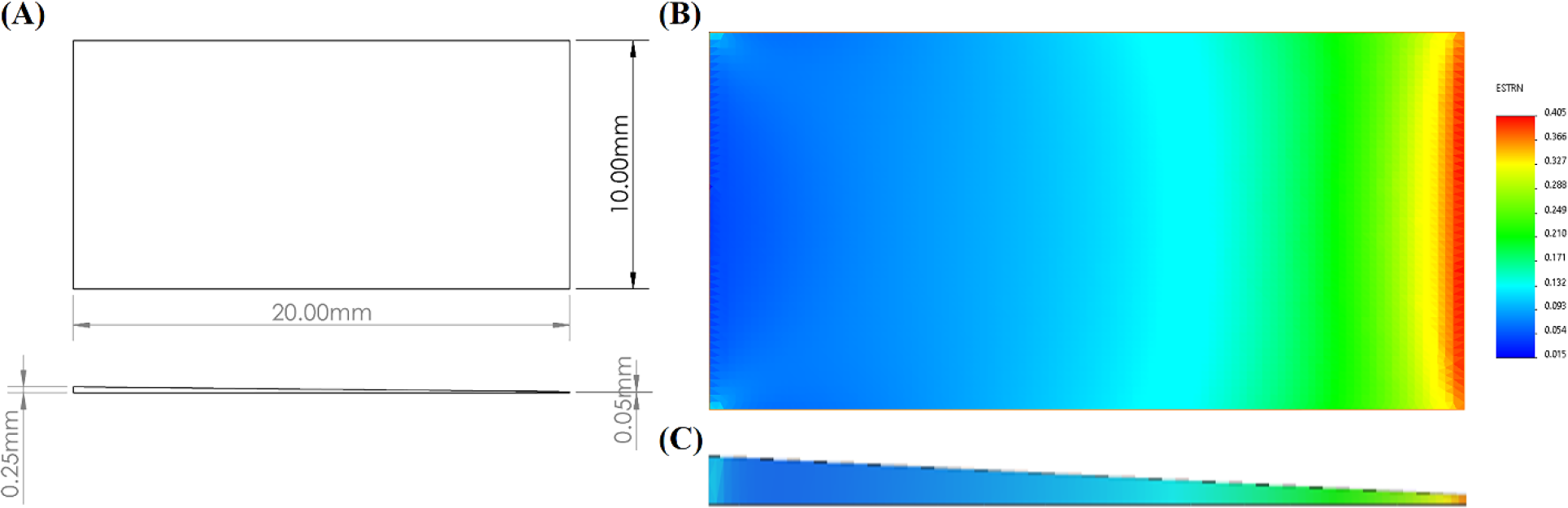
The SolidWorks simulation of the strain gradient. (A) The dimensions of the tapered membrane. (B) Top view and (C) cross-section of SolidWorks strain simulation.

While the computational simulation displayed the strain gradient, the presence of the gradient needed to be verified experimentally. To validate the actual strain at each of the previously mentioned points, ink dots were drawn on the membrane at 0.5 cm, 1.0 cm, and 1.5 cm from the thicker end, their deformations were recorded, and their percentage change in length was analyzed upon 15% overall stretch. As shown in Figure 5, the length of each dot along the stretch direction was increased, and then the strain was calculated in comparison with the original length before the stretching for each of the points. The actual strain at these location under 15% overall stretch was 9.8%, 16.0%, and 21.0% at 0.5 cm, 1.0 cm, and1.5 cm from the thicker end of the membrane, respectively (Table 2).

**Figure 5.**
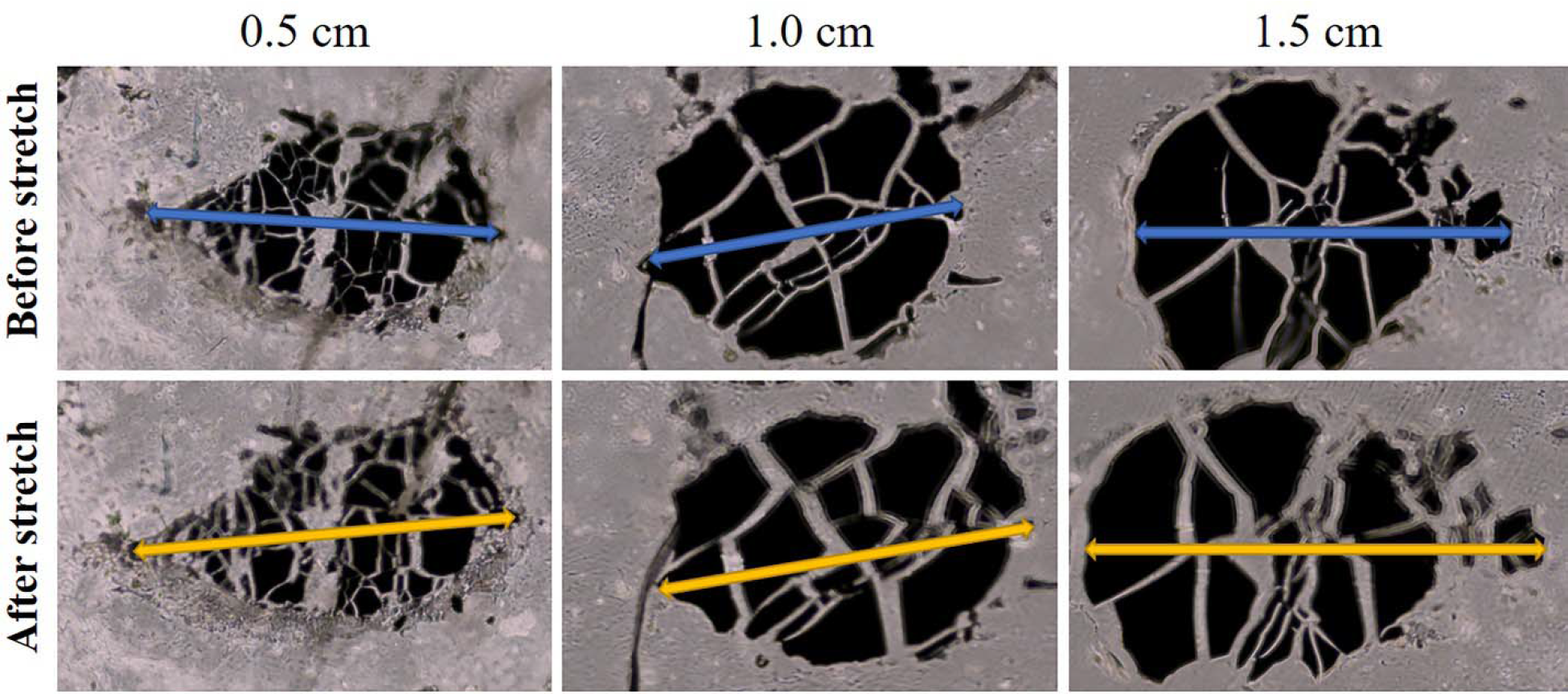
Optic images of marked dots on the membrane before and after exposed to 15% overall strain.

Table 2 compared the observed and simulated strains at different locations. Although there was difference between these values, which likely resulted from the approximation of the dot location in the experimental characterization, these sets of data revealed the formation of the strain gradient – smaller strain at a thicker membrane while larger strain at a thinner membrane.

### 3.3. Strain-dependent cell morphology

Under the static condition, the cells spread out in all directions and displayed random cell orientation. In comparison, depending on the magnitude of the mechanical strain, the cells could orient perpendicular to the stretching direction. As shown in Figure 6A, thicker regions of the PDMS membrane experienced smaller strains, resulting in less cell orientation in relation to the stretch direction. As the strain increased, the cells oriented themselves perpendicular to the stretch direction as seen through the nuclei and the F-actin. However, this trend diminished when the strain was further increased at the thinner regions of the PDMS membrane. Considering the difficulty in quantifying the orientation of cells, which displayed irregular morphology, the nuclei were more regular and easier to analyze. The orientation angle of the nucleus, as defined in Figure 6C, reflected this strain-dependent cell orientation as well. Ideally, under the static condition, the orientation of the cells and nuclei was completely random, and thus there should theoretically be 16.7% of nuclear (cell) population at each of the orientation angle ranges, or each of 6 sectors of 15° intervals (Figure 6B). Upon the mechanical stretching, at a smaller strain in region 1, nuclear orientation angles were mostly in the 30 - 90° range with a majority of cells in the 30 - 45° range. As the strain increased with a decrease in the membrane thickness in the regions 2 and 3, the endothelial cells displayed a stronger nuclear orientation perpendicular to the stretch direction. For example, in region 2 there was a greater percentage of nuclei with an orientation angle between 45° and 90° and most nuclei with an orientation angle between 75° and 90°, aligning more perpendicularly to the stretch direction. However, the trend shown in the regions 1, 2, and 3 became less significant in the regions 4 and 5. It was speculated that the regions 4 and 5 were overstretched, likely entering the pathogenic range of mechanical stretch (*e.g*., hypertension). At these regions, the cell and nuclear orientations became more randomly distributed, similar to the situation in region 1.

**Figure 6.**
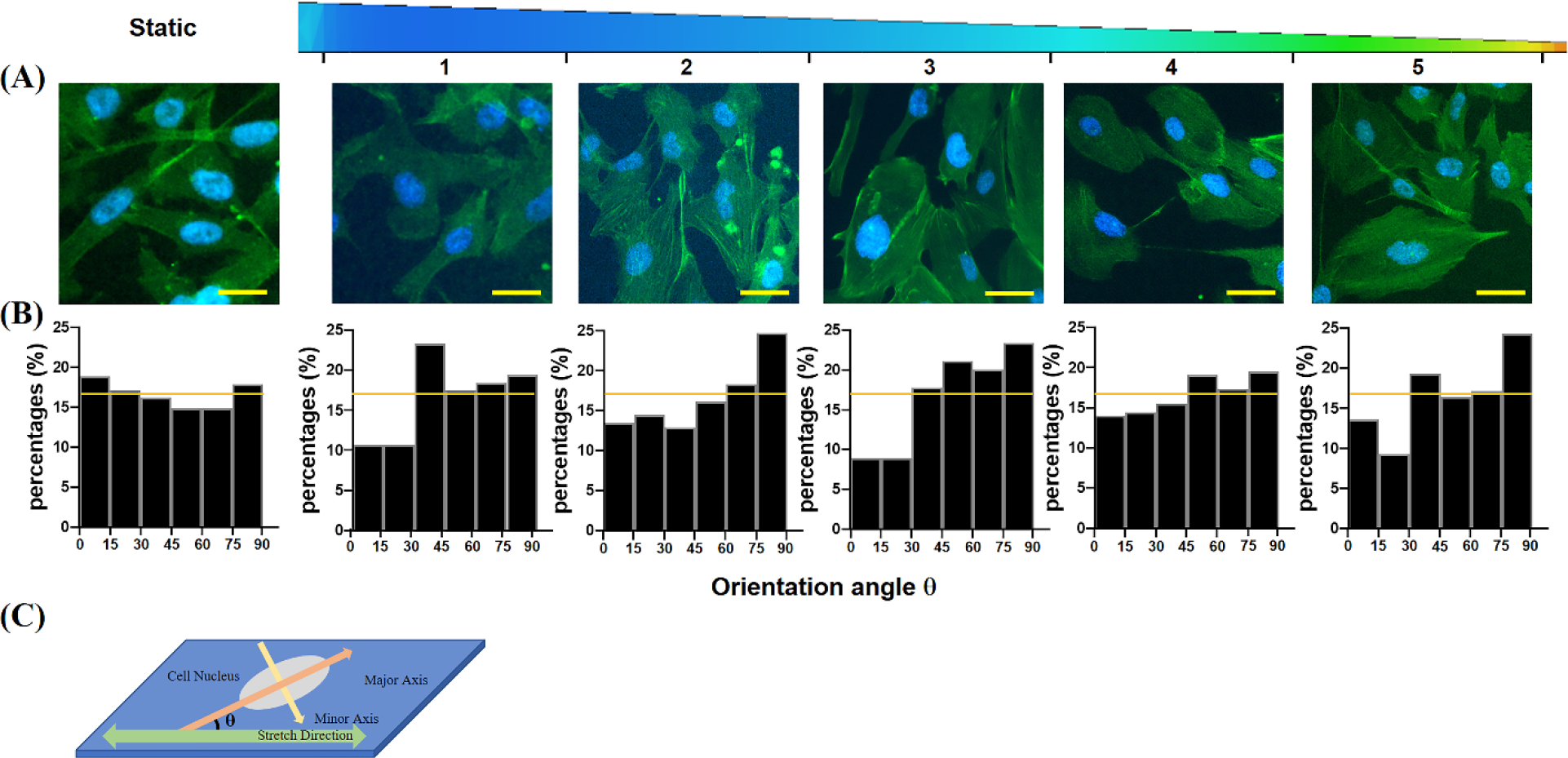
Mechanical strain-dependent cell/nuclear orientation. (A) Representative immunofluorescence images of hCMECs subjected to mechanical stretch of various strains in comparison with the static condition. The strain increases from region 1 to region 5. The nuclei are stained in blue and F-actin stained in green. Scale bars are 50 μm. (B) Quantitative analysis of nuclear orientation of hCMECs under the static condition and various strains. The horizontal lines represent the ideal nuclear orientation under static condition. (C) Illustrative definition of the nuclear orientation angle θ.

The nuclear elongation also showed the strain-dependent behavior. As shown in Figure 7, under the small strain in region 1, the nuclear elongation did not show a significant difference from the static condition. However, when the strain increased in region 2, the nuclei were stretched more than the static condition. When the strain was further increased from region 3 to 5, the overstretching led to deviation from the previously observed trend of, resulting in the decreased nuclear elongation from region 2.

**Figure 7.**
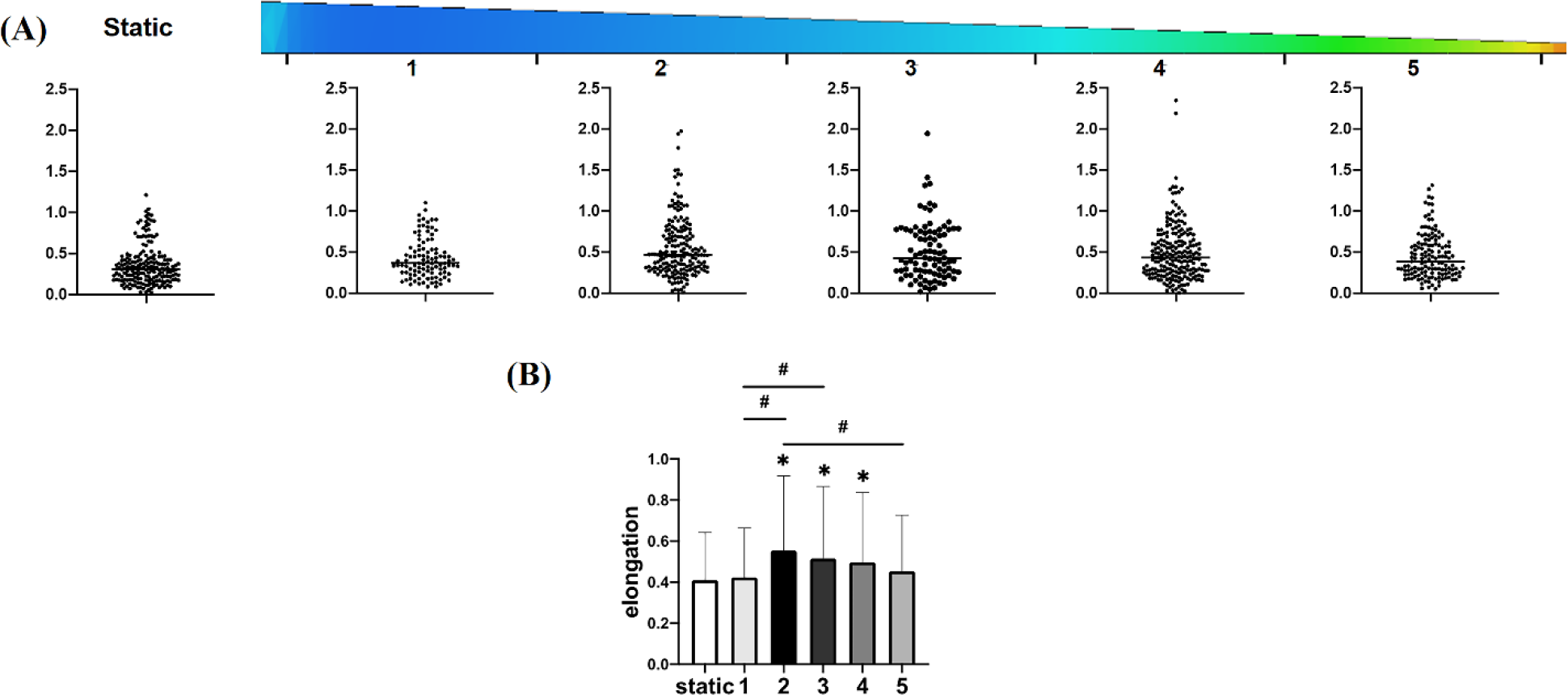
Strain-dependent nuclear elongation. (A) Dot plots of nuclear elongation of hCMECs under static condition and various mechanical strains. The strain increases from region 1 to region 5. (B) Comparison in nuclear orientation under static condition and various strains. *: *p* < 0.05 compared to the static controls. #: *p* < 0.05 between groups.

## 4. Discussion

This electromagnetically driven stretching device has precise control over the strain and frequency of the mechanical stretch. In addition to the gradient of strains ranging from 1.5% to 40% as demonstrated in this study, uniform strains can also be generated with a PDMS membrane of uniform thickness by altering the design of the track dimension (*i.e.*, the distance that the moving component is allowed to travel on the track). For example, when a PDMS membrane of 200 μm in thickness was used in the current design, a uniform strain of 14.8 ± 0.3% was generated on the membrane as characterized by the aforementioned dot deformation method, which agreed with the theoretical strain value of 15%. Furthermore, when 5 mm displacement is used, instead of 3 mm displacement in the current design, an overall 25% strain can be achieved. As such, this device spans the mechanical strain from physiological (5 – 10%) to pathological (>20% in hypertension) range. Moreover, the PWM signal generator has a frequency range between 1 Hz and 150 kHz, covering the physiological frequency range. Conversely, the reported electromagnetic stretching device generated the cyclic stretch at the strain of 1.4% and a frequency of 0.01 Hz, not in the physiological range^11^. Therefore, our electromagnetic stretching device can provide pathophysiologically relevant mechanical stretch. In addition, the use of the electromagnet separated from the container where the cell culture is conducted can effectively prevent the potential contamination from the stretching operation. It was noticed that the device ran out of battery after several hours of cyclic stretching. To sustain the cell stretching over days even weeks, a DC power source should be adopted to run the device continuously.

When the cell study was conducted on the strain gradient, the thinner end of the tapered PDMS membrane occasionally broke during the cyclic stretching because of the weakened mechanical strength. To resolve this problem, the membrane thickness needs to be increased. We therefore conducted the SolidWorks simulation with a thicker, tapered PDMS membrane. The results suggested that when the overall thickness was increased, a strain gradient could still be formed. For example, a strain gradient of 1.6% to 34% was generated on a tapered PDMS membrane with a thicker end at 400 μm and the thinner end at 100 μm. Although certain ranges of mechanical strains have been reported, these strain variations result from the edge effects and are heterogenous^5, 14^. A more controllable strain gradient has been demonstrated by applying geometrical constraints on a stretchable membrane. However, the strain gradient ranges between 12% to 18% across a distance of 1.5 mm, and the gradient is irregular^15^. Conversely, the strain gradients reported here are defined and can be tailored for the end application, and more importantly the mechanical strain/gradient can be generated over a large area, for instance, centimeter scale as demonstrated here.

The thicker and robust membrane allows us to add nanoscale structures on the top of the membrane to replicate the nanostructured extracellular matrix (ECM) and/or put soft hydrogel on the membrane to match the ECM stiffness. These nanostructures and soft hydrogel together with the pathophysiological mechanical stretch closely recapitulate the *in vivo* cell microenvironment.

Moreover, building this device is inexpensive. The 3-D printing of the stretching part costed the ink (~$2) and the printing charge (~$30). The electromagnet, function signal generator, and the switch costed $11, $16, and $1, respectively. These costs, together with other small parts such as wires, batteries, and PDMS membrane, were less than $70 in comparison with several thousand dollars for commercial Flexcell Systems. To sum up, this novel electromagnetic stretching device is simple, versatile, and cost-effective, suitable for the study of mechanical stretch effects.

Furthermore, the cell study exhibited that the microvascular endothelial cells responded differently to the alteration in mechanical strain. Physiologically relevant mechanical strain can effectively orient the endothelial cells perpendicular to the stretch direction while excessive strain may harm the endothelial phenotype. These observations agreed with the *in vivo* results. The functionality of tight junctions between adjacent endothelial cells is significant because they form an adjacent barrier between cells to prevent the leakage of molecules and ions through the plasma membranes of adjacent cells. Tight junctions can become impaired under pathological stress, leading to increased membrane permeability, or leaky endothelium. The ongoing research will examine how pathophysiological mechanical stretching affects the formation and function of vascular endothelium.

## 5. Conclusion

This novel electromagnetically driven 1-D stretching device provides a simple, versatile, cost-effective, pathophysiologically relevant platform that enables us to study the effects of mechanical strain, both normal and abnormal, on cells and tissues. The ability for this device to span the pathophysiological range allows us to observe the behavior of cells under hypertension and understand its effects on targeted cells, tissues, and organs contributing to various diseases. This device will also provide a useful *in vitro* platform for other biological studies involving mechanical stretching.

